# Phenotypic and transcriptomic responses of the sun- and shade-loving plants to sunlight and dim-light conditions

**DOI:** 10.1101/2022.01.28.477942

**Authors:** Yu-Xin Zhang, Yu-Qian Niu, Xin-Feng Wang, Zhen-Hui Wang, Meng-Li Wang, Ji Yang, Yu-Guo Wang, Wen-Ju Zhang, Zhi-Ping Song, Lin-Feng Li

## Abstract

Elucidating how plant species respond to variable light conditions is important to understanding the ecological adaptation to heterogeneous natural habitats. However, the phenotypic responses and gene regulatory network of shade-loving plants under distinct light conditions have remained largely unclear. In this study, we assessed the differences in phenotypic and transcriptomic responses between Arabidopsis (sun-loving) and *Panax ginseng* (shade-loving) to sunlight and dim-light conditions. Our results showed that, compared to the Arabidopsis, ginseng plants not only exhibited a lower degree of phenotypic plasticity in response to distinct light conditions, but also showed higher photosynthetic efficiency under dim-light conditions. Further time-course transcriptome profiling and gene structural analyses revealed that differentially transcriptional regulation together with increased copy number of the photosynthesis-related genes (*i.e*., electron transfer and carbon fixation) may improve the photosynthetic efficiency of ginseng plants under dim-light conditions. In contrast, the loss-function and inactivation of phytochrome-interacting factors are potentially associated with the observed low degree of phenotypic plasticity of ginseng plants under the changing light conditions. Our study provides new insights on how shade-loving plants respond to variable light conditions. Candidate genes related to shade adaptation in ginseng provide valuable genetic resources for future molecular breeding of high-density planting crops.

**Highlight:** The shade-loving species *Panax ginseng* possesses lower phenotypic plasticity under distinct light conditions and shows high photosynthesis efficiency under dim-light condition.

## Introduction

Elucidating the molecular mechanism underlying plant adaptation is one of the fundamental topics in evolutionary biology. As sessile organisms, plant species have evolved sophisticated strategies to respond to changing environmental conditions (Lee *et al*., 2007). As an essential environmental factor, sunlight not only provides source of solar energy for photosynthesis but also serves as an information signal in regulating plant growth and development (Hao *et al*., 2012; Paik *et al*., 2017). However, both the light quality and quantity vary spatially and temporally in natural habitats, with shaded environments imposing constraints on plant habitability due to light resource limitation (Cookson and Granier, 2006; Evers and Bastiaans, 2016). To survive, plant species have evolved two opposing strategies to deal with shaded environments, namely shade avoidance and shade tolerance (Gommers *et al*., 2013).

The shade-avoidance strategy refers to changes in growth characteristics under shaded environments, including elongation of seedling hypocotyls, stems and petiole elongation, branch number reduction and accelerated flowering date, collectively termed the shade-avoidance syndrome (SAS) (Ren and Gray, 2015). These shade-avoidance responses confer the ability of individual plants to escape shaded environments and complete their life cycle before the canopy becomes crowded (Callahan *et al*., 1997). Molecular mechanism underlying the SAS has been well documented in a variety of heliophytic (sun-loving) plants (Casal, 2013; Yang and Li, 2017). For example, evidence from the shade-intolerant species *Arabidopsis thaliana* (referred to as Arabidopsis) has clearly illustrated that shade-avoidance responses are regulated by genes involved in photoreceptor signaling networks, such as *PHYA/B* and *CRY1/2* (Hornitschek *et al*., 2012; Pedmale *et al*., 2016).

Plant species growing in canopy-shaded habitats adopt an alternative shade-tolerance strategy to respond to the dim-light environments (Jia *et al*., 2020). Shade tolerance is the ability of a given plant species to efficiently modify its morphology and physiology to cope permanently with shaded habitats (Valladares and Niinemets, 2008). Common phenotypes of the shade-tolerant plants include reduced chlorophyll a:b ratio, increased specific leaf area and photosystem (PS) II:I ratio (Gommers *et al*., 2013). However, molecular bases underpinning the shade-tolerance phenotype still remained under-investigated. To date, three hypotheses based on the photomorphogenesis regulatory network have been proposed, all of which invoke suppression of the SAS: differential control of phytochrome-interacting factors (PIFs), increased involvement of molecular shade-avoidance antagonists, and specific regulation of phytohormone biosynthesis (Gommers *et al*., 2013; Leivar *et al*., 2012). For example, upregulation of the *PHYA* gene acts as an antagonistic factor to inhibit hypocotyl elongation in both the sun-loving Arabidopsis and shade-loving *Cardamine hirsuta* (Molina-Contreras *et al*., 2019; Yang *et al*., 2018).

On the other hand, the ecological and physiological perspective holds that shade tolerance is not solely the suppression of SAS but also an evolutionary strategy of plant species to allocate energy and resource toward photosynthesis (growth) and physical defense (survival) (Valladares and Niinemets, 2007). Based on this assumption, two partly contrasting ecological strategies have been proposed to explain the survival-growth trade-off in shade-tolerant species: maximization of net carbon gain in low light (carbon gain hypothesis) and maximization of the resistance to biotic and abiotic stresses in the understory (stress tolerance hypothesis) (Givnish, 1988; Kitajima, 1994; Valladares and Niinemets, 2007). The carbon gain hypothesis defines shade tolerance as the maximization of photosynthetic carbon gain in low light but with the minimization of respiration costs for maintenance (Givnish, 1988). Under this hypothesis, any morphological and physiological traits that improve light capture and use efficiency are considered advantageous for increasing plant shade tolerance. Three mechanisms related to photosynthesis efficiency have been characterized(Kaiser *et al*., 2019): (*i*) the acceleration of non-photochemical quenching (NPQ) relaxation upon shift to low light (Miyake *et al*., 2005), where molecular bases of NPQ relaxation involve photosynthesis-related genes (*i.e*., light harvesting antenna proteins) and their regulatory genes (*i.e*., *KEA3* and *ZEP*) (Armbruster *et al*., 2016); (*ii*) the activation/deactivation of genes involved in Calvin-Benson-Basham cycle (*i.e*., *rbcL* and *RCA*) (Chen *et al*., 2015); and (*iii*) the alteration of stomatal conductance, where genes involved in stomatal dynamics during light fluctuations include blue- and red-light regulatory genes (*i.e*., *Phot1* and *Phot2*) (Hosotani *et al*., 2021). Conversely, the stress tolerance hypothesis argues that survival in shaded habitats is determined by the ability of plants to resist biotic and abiotic stresses rather than to maximize carbon gain (Kitajima, 1994; Poorter and Kitajima, 2007). Shade-tolerant plants usually are expected to show higher longevity of their organs at low daily light integral by increasing the ability to guard against mechanical damage, herbivores and pathogens (Cook-Patton *et al*., 2020). Together, these empirical studies suggest that shade tolerance is a complicated adaptive strategy that not only relies on diverse morphological and physiological characteristics but also differs obviously among species in natural habitats. With this reasoning, it is of particularly important to expand the investigations with more species and multi-disciplinary approaches.

In this study, we aimed to address whether *Panax ginseng* C.A. Meyer (referred to as ginseng) has evolved specialized phenotypes to cope with the shaded habitats where it naturally grows. Then, we next elucidated the molecular bases underpin this ecological adaptation. Ginseng is a perennial herbaceous species within the genus *Panax* of the family Araliaceae (Valcarcel *et al*., 2017; Wen and Zimmer, 1996). As a highly valued and popular medicinal plant, extensive studies have been made on its evolutionary history and pharmacological effects (Li *et al*., 2017; Shi *et al*., 2015). It has been documented that ginseng is a typical sciophytic (shade-loving) species showing photo-inhibitory symptoms under intense light conditions (> 500 μmol·m^-2^·s^-1^) (Jung *et al*., 2020; Miskell *et al*., 2002). This attribute makes ginseng an ideal system to address its phenotypic and molecular responses to variable light conditions. With this reasoning, we performed detailed phenotypic comparisons between ginseng and the heliophytic model plant Arabidopsis under simulated sunlight, shade and deep-shade light conditions. To further address the underlying genetic bases and transcriptomic responses, we examined the time-course expression profiling of ginseng under the three simulated light conditions. The aims of our study were to: (1) address whether ginseng has evolved specific morphological and physiological traits to cope with shaded habitats; (2) elucidate the molecular bases underlying the specialized phenotypes. As energetic investment in the SAS is a major contributor to crop yield loss, candidate genes identified in shade-tolerant species can provide a novel avenue to breed for crops that are tolerant of high-density planting conditions.

## Materials and Methods

### Plant materials and cultivation conditions

The typical heliophytic (Arabidopsis) and sciophytic species (ginseng) were used to evaluate phenotypic plasticity under simulated sunlight and dim-light (shade and deep-shade) conditions. The Columbia ecotype of Arabidopsis were used to measure the phenotypic and transcriptomic responses to distinct light conditions in this study. As ginseng is a perennial herb, we chosen both the one-year and four-year plants to measure the morphological and physiological traits. To examine how the same individual responds to changing light conditions, we only examined the transcriptomic response of the four-year ginseng plant. All the ginseng plants were collected from Jilin Province of China. For the phenotypic experiments, geminated seeds of Arabidopsis and ginseng (one-year) were transferred to the three independent light incubators (Ningbo jiangnan GXZ-280C, Ningbo, China). The four-year ginseng plants were also moved to the same light incubators when the leaf buds are sprouted. Light quantity and quality of the three incubators were set according to the spectral composition of light in natural habitats (Table S1) and previous studies (Molina-Contreras *et al*., 2019; Yang *et al*., 2018). The three simulated light conditions were: (1) sunlight, with red light/far-red light (R:FR) ratio = 12, light intensity = 280 μmol·m^-2^·s^-1^; (2) shade, R:FR = 0.4, light intensity = 30 μmol·m^-2^·s^-1^; and (3) deep-shade, R:FR = 0.2, light intensity = 20 μmol·m^-2^·s^-1^ (Fig. 1A). Far light is provided by the light-emitting diode lamp (KOUGIN, ShanDong, China). The Plant Lighting Analyzer (PLA-30 V2.00, Hangzhou, China) was employed to measure the light conditions. All experiments were performed under long day conditions (16 hours light/8 hours dark at 18°C).

**Figure 1.**
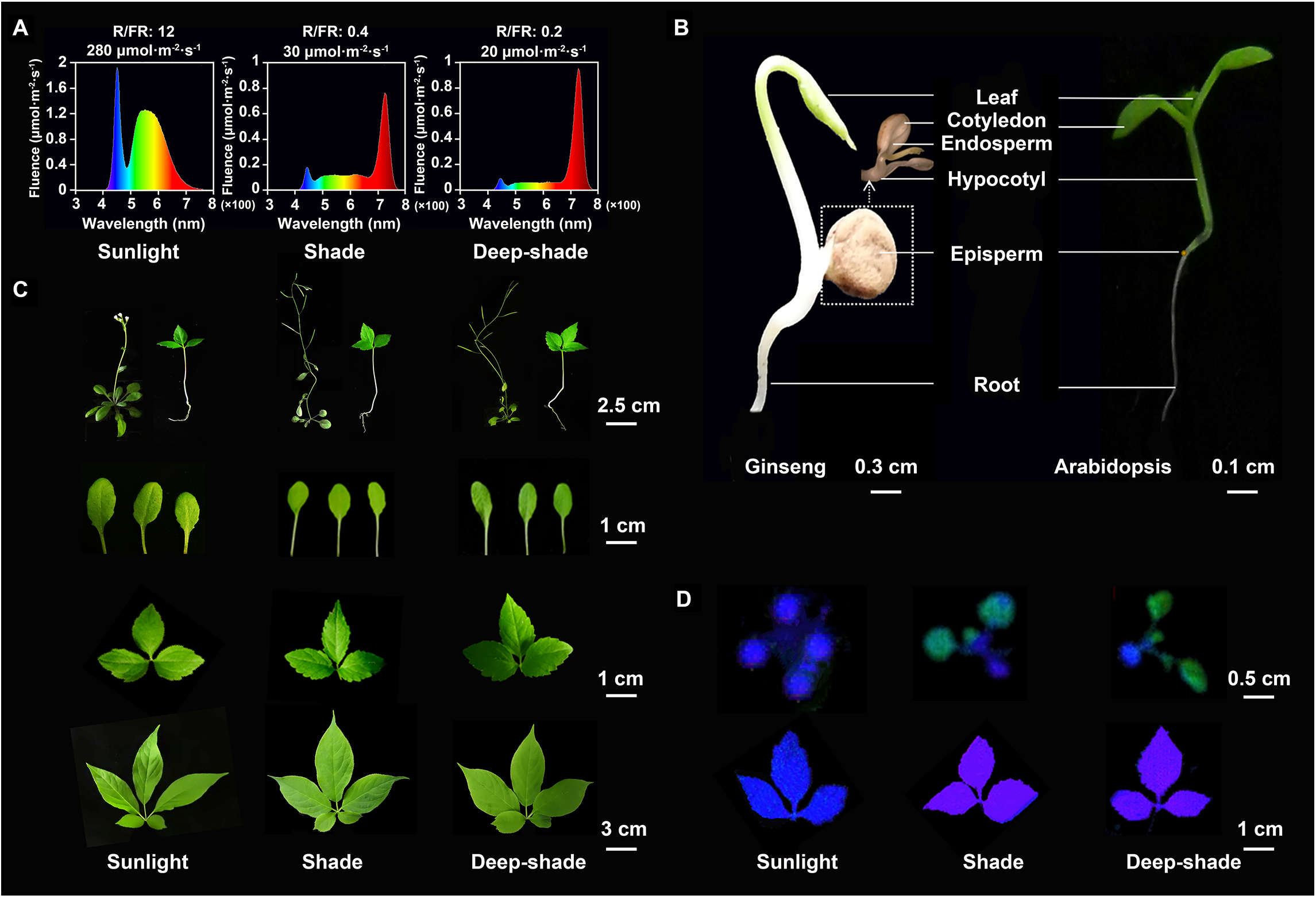
Phenotypic responses of Arabidopsis and ginseng to simulated sunlight and dim-light conditions. (A) Spectral composition of the three simulated light conditions. The color scheme indicates different light components in each simulated condition. Numbers on the top of each subpanel are the photosynthetic photon flux density (PPFD) and red (R)/far red (FR) ratio for sunlight (280 μmol·m^-2^·s^-1^, R/FR=12), shade (30 μmol·m^-2^·s^-1^, R/FR=0.4) and deep-shade (20 μmol·m^-2^·s^-1^, R/FR=0.2) light conditions. X- and Y-axis are the light wavelength and fluence, respectively. (B) Seedling morphologies of ginseng (left, ten days) and Arabidopsis (right, seven days). The white lines indicate the tissues of the two species. (C) Morphological traits in Arabidopsis and ginseng under sunlight (left), shade (middle) and deep-shade (right) light conditions. The first row is of the Arabidopsis (left, four weeks) and one-year ginseng (right, four weeks) plants under the three light conditions. From the second to fourth rows are the leaf morphologies of Arabidopsis (2^rd^ row), one-year (3^rd^ row) and four-year (4^th^ row) ginseng plants. (D) Chlorophyll fluorescence (Fv/Fm) imaging of Arabidopsis (upper row) and ginseng (lower row) under the three light conditions. Colors from green to blue and purple indicate low, median and high Fv/Fm signal intensities. Scale bar in each subpanel indicates the plant size.

### Measurement of morphological and physiological traits

To compare the phenotypic responses of the two species to distinct light conditions, both the Arabidopsis and ginseng plants grown in the three light incubators for three weeks were selected to conduct phenotypic measurements. Morphological traits of the leaf area and petiole length were photographed using a Nikon COOLSCOPE II (Nikon Corporation, Tokyo, Japan) and measured using the software ImageJ (Zhou *et al*., 2018). Flowering time of Arabidopsis and four-year ginseng plants were estimated based on the number of days from germination to the emergence of first flower. As the ginseng plants typically flower at the third year, the one-year ginseng plants were not included in the statistic of flowering time. In addition, we also measured several light-response physiological traits of the two species, including leaf anatomical structure, chlorophyll content and photosynthetic parameters. For the leaf anatomical structure, 45 and 90 paraffin sectioning samples were obtained from the leaf tissue of Arabidopsis and ginseng, respectively. Fresh leaf tissue was fixed with formalin-acetic acid-alcohol and embedded in paraffin wax. Paraffin-embedded sections were cut and stained with safranin-O and fast green. Based on these paraffin sectioning samples, we estimated leaf thickness using a Nikon Eclipse E200 optical microscope (Nikon Corporation, Tokyo, Japan). Chlorophyll a and b were extracted using an acetone and alcohol protocol(Zhao *et al*., 2018) and quantified using Synergy™2 multi-detection microplate reader (Biotek, VT). Light response curve was measured based on the red and blue light sources using Li 6800 (Li-COR, Lincoln, USA). Maximum quantum yield of PSII (Fv/Fm) was calculated for Arabidopsis and ginseng plants using the program ImagingWin on plant phenotype imaging platform (Taiwan, China).

### Sampling, RNA sequencing and gene expression profiling

To address the transcriptomic responses of ginseng and Arabidopsis under the above three light conditions, all plants of the two species were grown under their simulated natural habitats. In brief, the four-year ginseng plants were grown under shaded light condition in greenhouse (at 18°C) at Fudan University in Shanghai (China). The Arabidopsis plants were grown under simulated sunlight condition (at 18°C, light intensity = 110 μmol·m^-2^·s^-1^, 16/8 hour) in light incubator (Ningbo Jiangnan GXZ-280C, Ningbo, China). All the Arabidopsis and ginseng plants were homogenized at low intensity sunlight (R:FR = 12, 110 μmol·m^-2^·s^-1^) for six hours. Then, we transferred these homogenized plants to the three light incubators corresponding to simulated high intensity sunlight (R:FR = 12, 280 μmol·m^-2^·s^-1^), shade (R:FR = 0.4, 30 μmol·m^-2^·s^-1^) and deep-shade (R:FR = 0.2, 20 μmol·m^-2^·s^-1^) light conditions. Gene expression profiling of four-year ginseng plants were surveyed at seven time-points (0 h, 0.25 h, 0.5 h, 1 h, 3 h, 6 h and 12 h). The molecular response and regulatory network for the SAS and photosynthetic efficiency have been well-documented in Arabidopsis (Chen and Chory, 2011; Franklin and Quail, 2010; Lin, 2002). This study only sampled four time-points (0 h, 1 h, 3 h and 6 h) to confirm its molecular responses under simulated sunlight and dim-light conditions. Leaf samples of the four-year ginseng plant were collected from the same individual at seven time-points mentioned above. Leaf samples of the Arabidopsis were collected from different individuals because of its small plant size. Leaf samples collected at 0 h under simulated sunlight (R:FR = 12, 60 μmol·m^-2^·s^-1^) were served as control. Each of the three light conditions contained three biological replicates.

All fresh leaf samples were immediately frozen in liquid nitrogen and stored at −80°C. Total RNA was extracted from each sample using RNA extraction kits (TIANGEN, Beijing, China) and quantified with Nanodrop 2000C spectrophotometry (Thermo scientific, Waltham, USA). Transcriptome sequencing was performed using Illumina Novaseq 6000 platform (Illumina, San Diego, USA). Clean data were obtained by removing the short reads containing adapter sequences, poly-N, and low-quality reads (Q value < 20). Clean reads were mapped onto the respective reference genomes using HISAT2 (Kim *et al*., 2019). Raw mapped read counts were calculated using the prepDE.py script providing by StringTie (Pertea *et al*., 2015). Differences in transcription level of each gene were estimated using DESeq2 (Love *et al*., 2014). Differentially expressed genes (DEGs) was defined according to the two-fold change differences (*p* < 0.05) in transcription level between the treatment (0.25-12 h) and control (0 h) time-points. Relative transcription level of the photosynthesis-related genes was also measured between the shade and sunlight conditions at the same time-point. Gene expression-level diversity and specificity were calculated according to previous study (Martinez and Reyes-Valdes, 2008).

### Identification of the candidate genes

The photomorphogenesis and photosynthesis genes in Arabidopsis were obtained from previous studies (Casal, 2012; Kanehisa and Goto, 2000; Martinez-Garcia *et al*., 2014) and Kyoto Encyclopedia of Genes and Genomes (KEGG) database (Kanehisa and Goto, 2000). Arabidopsis genome annotations (TAIR10) were obtained from TAIR (www.arabidopsis.org). Orthologous genes in ginseng were identified by searching the protein sequences of Arabidopsis against the ginseng genome using BLASTP (Altschul *et al*., 1990), with e-value 10^-5^ and 40% identity cutoff. Gene structure of the *PHYB* and *PIFs* genes was inferred according to previous study (Li *et al*., 2011). The homology model of *PHYB* gene was built by the SWISS-MODEL server (Waterhouse *et al*., 2018). Functional annotation of the ginseng genes was performed based on the KEGG and Gene Ontology (GO) databases (Harris *et al*., 2004; Kanehisa and Goto, 2000). Functional enrichments of the DEGs were carried out with the R package ClusterProfiler (Yu *et al*., 2012). A Venn diagram of expression overlap under different light conditions was drawn by VennDiagram (Chen and Boutros, 2011). A heatmap of the DEGs was plotted using pheatmap (version 1.0.12). WGCNA was performed for both ginseng and Arabidopsis using the WGCNA R package (Langfelder and Horvath, 2008).

## Results

### Phenotypic responses of Arabidopsis and ginseng under distinct light conditions

We assessed the differences in phenotypic responses between Arabidopsis and ginseng under simulated sunlight and dim-light conditions (Fig. 1A). Arabidopsis is a typical heliophytic species whose cotyledons often grow above the ground by elongating the hypocotyl during the seed germination (Fig. 1B). Thus, Arabidopsis seedlings can convert solar energy to chemical energy through photosynthesis at cotyledons. In contrast, the ginseng has hypogeal gemination with hypocotyls that lack the ability to elongate; as such, the young seedling relies solely on the endosperm for its required energy. In addition, Arabidopsis plants showed a higher degree of phenotypic plasticity in leaf morphology and anatomical structure, whereas both the one-year and four-year ginseng plants maintained stable phenotypes under the three light conditions (Fig. 1C, Supporting Information Fig. S1 and Table S2). Furthermore, Arabidopsis plants exhibited significantly lower maximum quantum yield of PS II at dim-light (Fv/Fm = 0.56-0.59) compared to sunlight (Fv/Fm = 0.84, *p* < 0.01) condition (Fig. 1D and Table S3). In contrast, the ginseng plants possessed higher maximum quantum yield at dim-light (Fv/Fm = 0.81) relative to sunlight (Fv/Fm = 0.75, *p* < 0.01) condition.

### Distinct levels of the phenotypic plasticity between ginseng and Arabidopsis

We compared photosynthesis- and reproduction-related traits between Arabidopsis and ginseng to identify the shade-tolerance phenotypes. Firstly, we measured the light response curve of both Arabidopsis and ginseng plants under sunlight condition. The heliophytic species Arabidopsis showed a markedly higher light saturation point (700-800 μmol·m^-2^·s^-1^) relative to the sciophytic species ginseng (300-400 μmol·m^-2^·s^-1^) (Fig. 2A and Table S4). We next reexamined specific morphological and physiological traits to confirm the above-mentioned phenotypic plasticity (see Fig. 1C). The Arabidopsis plants showed typical SAS under dim-light (shade and deep-shade) condition, including accelerated flowering, increased petiole length and decreased leaf area/thickness (Fig. 2B-F, Fig. S1 and Table S2). In contrast, both the one-year and four-year ginseng plants possessed apparent shade-tolerance phenotype under the dim-light conditions, such as increased leaf area but with relatively stable flowering time and petiole length (Fig. 2B-F, Fig. S1 and Table S2). It is notable that while both Arabidopsis and ginseng showed increased total chlorophyll content under dim-light conditions, they possessed distinct variability in chlorophyll a and b compositions (Table S2). Overall, the Arabidopsis plants showed relatively higher degree of decreased chlorophyll a/b ratio compared to ginseng under dim-light conditions (Fig. 2F).

**Figure 2.**
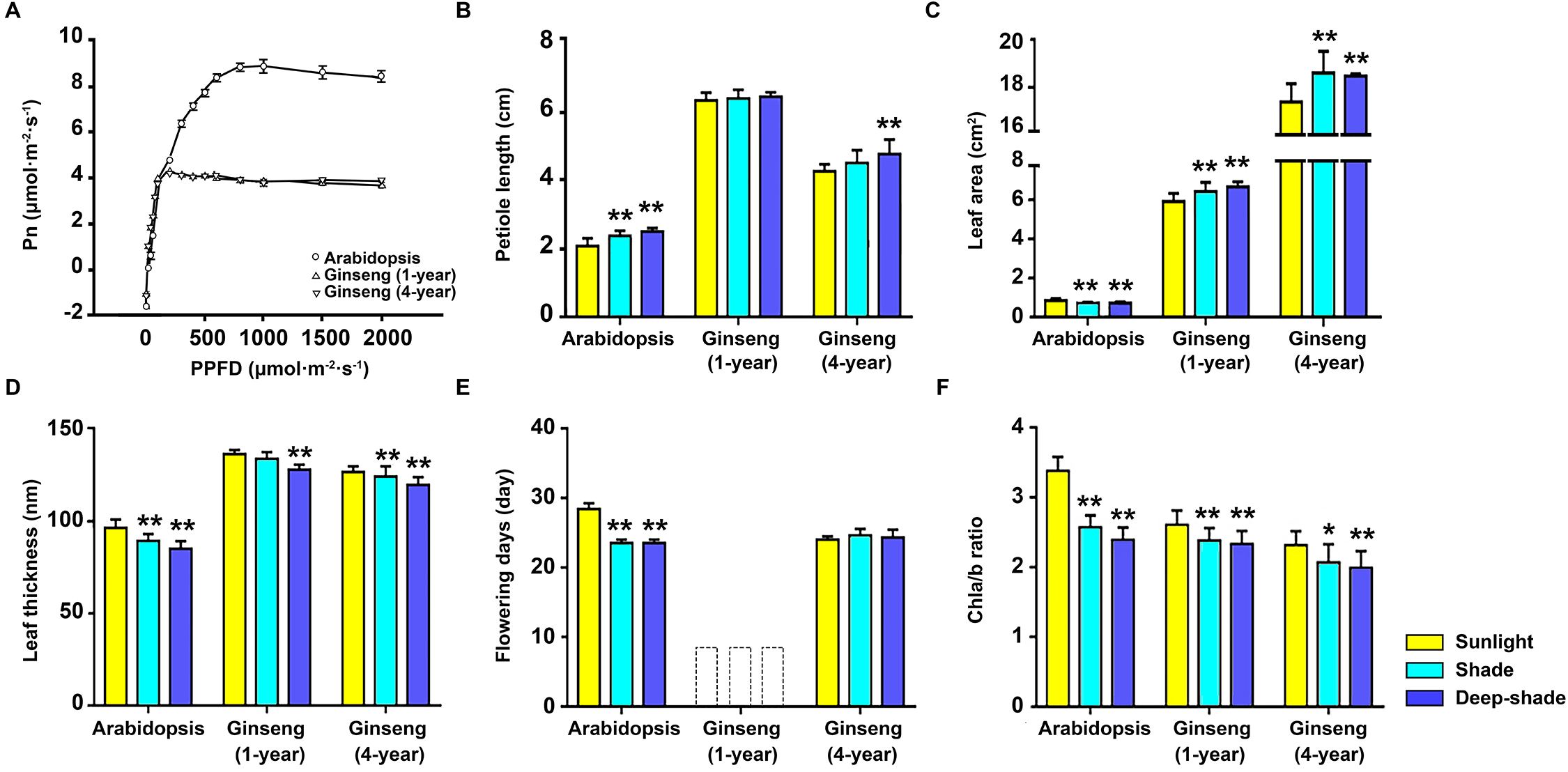
Phenotypic differences between the Arabidopsis and ginseng under sunlight and dim-light conditions. (A) Light response curves for Arabidopsis and ginseng. X- and Y-axis are net photosynthetic rate (Pn) and PPFD. (B-F) Statistics of the petiole length, leaf area, leaf thickness, flowering days, chlorophyll a/b ratio under the three light conditions. The ginseng plants usual flower after two years. Flowering time of the one-year ginseng plant was not calculated (dotted box in E). Significant test was performed by comparing the two dim-light conditions with sunlight condition. The data represent the means ± s.e. (n ≥ 10). (One-way ANOVA test, \**p* < 0.05; \*\**p* < 0.01).

### Overall gene expression patterns under three simulated light conditions

The above phenotypic comparisons revealed high adaptability of the ginseng plants to dim-light conditions compared to Arabidopsis. We thus analyzed time-course gene expression profiling to elucidate the underlying molecular responses. At the whole genome level, overall gene expression pattern of the ginseng plant under sunlight condition differed obviously from those expressed under the two dim-light conditions. For example, while the expressed-genes possessed similar expression-level diversity between the sunlight (*Hj* = 12.96-13.36) and dim-light (*Hj* = 12.84-13.54) conditions, they showed apparently different expression specificity between sunlight (*δj* = 0.05-0.08) and dim-light (*δj* = 0.02-0.06) conditions (Fig. 3A). Distinct overall expression patterns between the sunlight and dim-light conditions were also observed at the subgenome level (Fig. S2A-B). However, expressed-genes under sunlight condition exhibited less variability in terms of the *Hj* and *δj* values. In contrast, overall expression pattern of the Arabidopsis exhibited similar divergent among the three light conditions (Fig. 3A).

**Figure 3.**
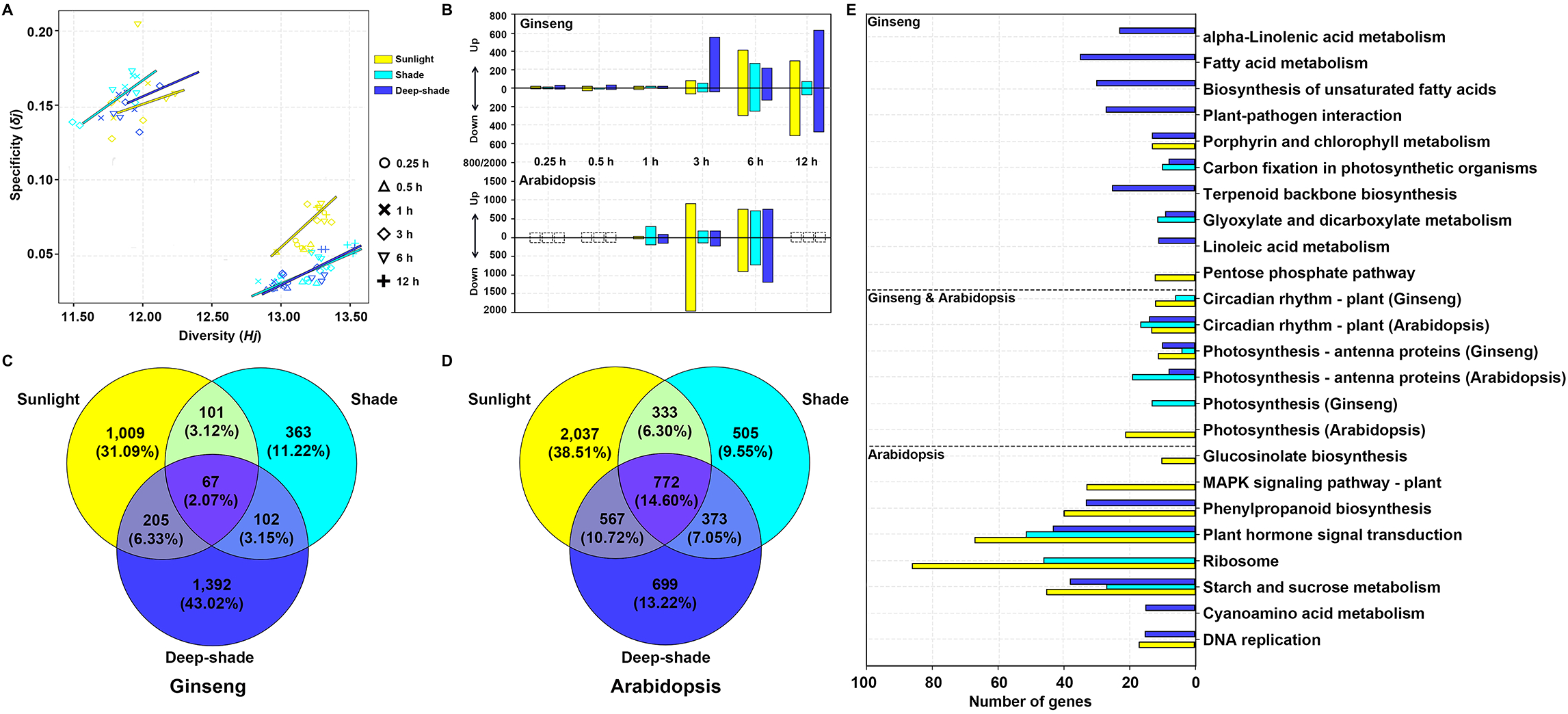
Overall gene expression pattern of ginseng and Arabidopsis leaf tissue under sunlight and dim-light conditions. (A) Visualizing transcriptome data based on expression diversity (*Hj*) and specificity (*δj*). Each symbol represents a sampling time point. The colored lines on up-left and down-right corners are Arabidopsis and ginseng, respectively. (B) Number of differentially expressed genes (DEGs) under sunlight and dim-light conditions in ginseng (top) and Arabidopsis (bottom). The up- and down-regulation were defined according to the 2-fold change of expression level between the treatment (0.25-12 hour) and control (0 hour) time point. Dashed boxes in the bottom subpanel indicate samples were not collected at these time-points in Arabidopsis. (C-D) Venn diagrams of the DEGs under sunlight and dim-light conditions in ginseng and Arabidopsis. Numbers within each circle were obtained from the comparisons between the control and six treatment time points. (E) Enriched KEGG pathways of the DEGs under sunlight and dim-light conditions. Top middle and bottom subpanels represent the ginseng-specific, ginseng and Arabidopsis shared and Arabidopsis-specific KEGG pathways.

It is notable that gene-expression dynamics of the four-year ginseng differed obviously at the 3-hour time-point under the three light conditions (Fig. 3A and Fig. S2 A-B). This pattern was also confirmed by the overall expression distance where samples before the 3-hour time-point were clustered as a clade (excepting the 0.25 h under deep-shade) (Fig. S3A). In line with this observation, numbers of DEGs also increased dramatically from the 3-hour time-points (Fig. 3B). In particular, up- and down-regulated genes exhibited similar expression pattern after 3-hour under all the three light conditions (Fig. S4A). Likewise, the number of DEGs and overall expression distance among the time-points also increased obviously in the Arabidopsis after the hour time-point (Fig. 3B and Fig. S3B). These findings suggest that most of the genes involved in the response to light condition were activated at the 3-hour in leaf tissue of the ginseng and Arabidopsis.

Intersection analysis of the overall DEGs revealed that a total of 363-1,392 (11.22-43.02%) and 505-2,037 (9.55-38.51%) of the DEGs were specific to the three light conditions in ginseng and Arabidopsis, respectively (Fig. 3C-D). By contrast, only 67-205 (1.79-6.56%) and 333-772 (6.30-14.60%) of the DEGs were commonly identified in two or all three of the light conditions in the two species. In the two subgenomes of the ginseng, we also observed similar proportions of the specific (11.46-43.10% and 11.05-43.31%) and common DEGs (1.82-6.66% and 2.33-6.10%) DEGs (Fig. S5). KEGG enrichment analysis of the DEGs identified 11 and 13 significantly enriched pathways under the three light conditions in Arabidopsis and ginseng, respectively (Fig. 3E and Table S4). In particular, the three light-response pathways (photosynthesis-antenna proteins, photosynthesis and circadian rhythm-plant) were commonly identified in the two species (Fig. 3E and Table S5). In line with this, GO analysis of the DEGs also identified several enriched terms related to photosynthesis and phytohormones, including light harvesting, chlorophyll biosynthetic process, response to blue/far red light and circadian rhythm (Table S6).

### Identification of candidate genes related to photosynthetic efficiency

The above analyses of time-course gene expression revealed that genes involved in photosynthesis were differentially expressed in the leaf tissue of both ginseng and Arabidopsis. We thus compared how these photosynthesis-related genes respond to distinct light conditions in the sun-loving (Arabidopsis) and shade-loving (ginseng) species. We firstly focused on the genes involved in chlorophyll a and b biosynthesis. In ginseng, a general pattern was that the majority of these genes showed immediately upregulation (but followed by down-regulation) in the two dim-light conditions (Fig. 4A and Table S7). In contrast, these genes exhibited a down-regulation pattern in across all the time-points in the sunlight condition. By comparing the relative expression level at the same time-point among the three light conditions, we found that five genes were significantly up-regulated under dim-light compared to sunlight condition (Table S8). For example, both copies of *CAO*, a key gene of the chlorophyll b synthesis, were significantly up-regulated under dim-light conditions. In Arabidopsis, most of these genes were down-regulated across the time-points in the three light conditions (Fig. S6 and Table S9). Notably, we identified four significantly up-regulation genes (*HEMA1, CHLH, CRD* and *CAO*) in the two dim-light conditions, with the *CAO* and *HEMA1* also up-regulated (but not significant) in the sunlight condition. This observation was further confirmed by the comparisons of relative gene expression level where most of these genes showed higher transcription level in the two dim-light compared to the sunlight condition (Table S10). These molecular responses may explain why the ginseng and Arabidopsis plants grown under dim-light conditions contained higher level of chlorophyll content compared to sunlight condition (see in Table S2).

**Figure 4.**
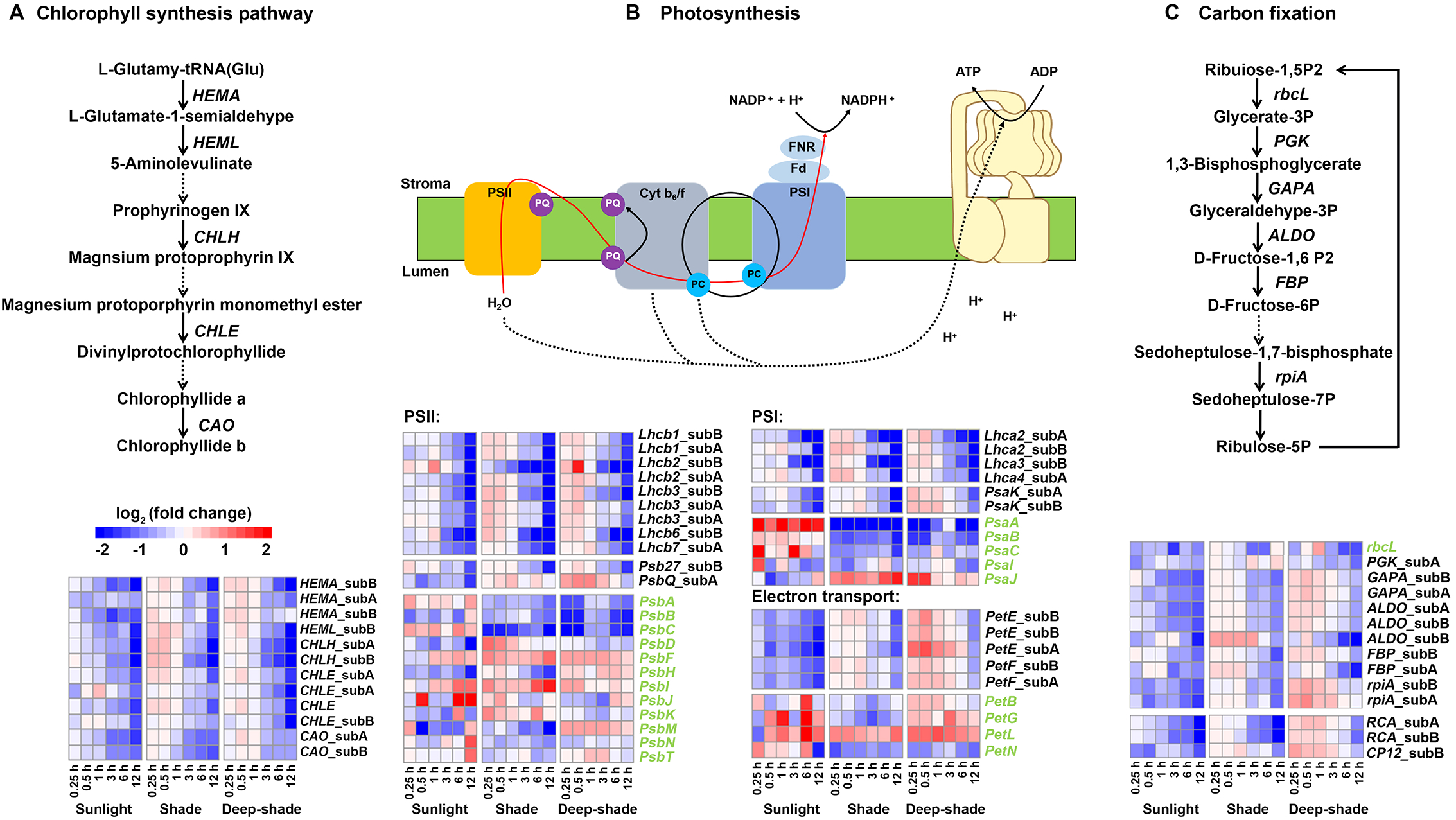
Expression pattern of the photosynthesis-related genes in ginseng leaf tissue under sunlight and dim-light conditions. (A) Differentially expressed genes (DEGs) involved in chlorophyll synthesis pathway. (B) DEGs involved in PSI, PSII and electron transport chain. (C) DEGs involved in carbon fixation. Gene names marked in black and green color represent nuclear and chloroplast genes, respectively. Color scheme indicates the up (red) and down (blue) of the genes between the treatment (0.25-12 h) and control (0 h) time point. “subA” and “subB” are orthologous genes from the two *Panax ginseng* subgenomes. PC: Plastocyanin; PQ: Plastoquinone; Fd: Ferredoxin; FNR: Ferredoxin-NADP+-oxidoreductase; Cyt b6/f: Cytochrome b6/f complex; PSI: Photosystem I; PSII: Photosystem II.

Both chlorophyll a and b interact with light-harvesting complex (LHC) proteins which convert solar energy into chemical energy in the chloroplast (Liu *et al*., 2013). Consistent with the expression patterns of chlorophyll synthesis genes, the majority of LHC-I and LHC-II genes showed up-regulation in ginseng leaf tissue under dim-light conditions (Fig. 4B and Table S7). In Arabidopsis, most of these light-harvesting genes also showed higher transcription level in the dim-light compared to sunlight condition (Fig. S6, Table S9-S10). However, it is unexpected that almost all of these genes were significantly down-regulated in the sunlight condition. Similar pattern was also observed in the nuclear encoding genes of PSI (*i.e., PsaD* and *PsaE*) and PSII (*i.e., PsbP* and *Psb28*) (Fig. S6 and Table S9-10). In contrast, opposite expression pattern was observed in these genes of PSI (*PsaA, PsaB* and *PsaC*) and PSII (*PsbA* and *PsbC*) in ginseng leaf tissue (Fig. 4B, Table S7-8). For example, transcription level of the *PsbA* gene, encoding the photosystem II response center protein D1, was significantly increased in ginseng leaf tissue under sunlight compared to dim-light conditions. However, no significant up-regulation of the *PsbA* gene was observed in the Arabidopsis leaf tissue. Different regulation pattern between the sunlight and dim-light conditions was also observed at the *PsaA* gene of ginseng and Arabidopsis leaf tissue. The electron mediators in the photosynthetic chain consist of plastoquinone (PQ), cytochrome (Cyt) b_6_/f complex, ferredoxin (Fd) and plastocyanin (PC). In ginseng, our results showed that these genes involved in photosynthetic electron transport chain are differentially expressed under the three light conditions (Fig. 4B and Table S7). For example, both the *PetE* and *PetF* gene, encoding PC and Fd, were significantly up-regulated under dim-light compared to sunlight condition. In contrast, the Cyt-b_6_/f complex gene *PetN* was up-regulated under sunlight condition. However, we did not observe significant up-regulation of these genes in Arabidopsis leaf tissue (Fig. S6, Table S9-10).

The photosynthesis system converts solar energy to NADPH and ATP, which provide chemical energy to fix carbon during the Benson-Calvin cycle. We therefore examined whether genes involved in carbon fixation showed differential expression under the three light conditions. In ginseng, our results showed that majority of the carbon fixation genes showed relative up-regulation under dim-light compared to sunlight condition (Fig. 4C, Table S7-8). However, most of these genes possessed similar expression patterns under the three light conditions in Arabidopsis (Fig. S6, Table S9-10). In ginseng, for example, both two copies of the *RCA* gene encoding Rubisco activase were also up-regulated under dim-light conditions (Fig. 4C, Table S7-8). In Arabidopsis, however, significant up-regulation of the *RCA* gene was observed in both the sunlight and dim-light conditions (Fig. S6 and Table S9). Together, these above findings suggest that transcriptional patterns of these genes are potentially associated with the distinct photosynthetic efficiency in the Arabidopsis and ginseng under different light conditions.

### Gene expression pattern of the photomorphogenesis regulatory network

The above morphological comparisons revealed different individual performance between the ginseng and Arabidopsis under sunlight and dim-light conditions (see Fig. 1 and 2). We thus examined how these photomorphogenesis regulatory genes were expressed in the leaf tissue of the two species under the three light conditions. Among the phytochromes and cryptochromes, only red-light photoreceptor *PHYB* showed significant up-regulation under dim-light conditions in both the Arabidopsis and ginseng (Fig. 5). The activated phytochromes (*PHYA* and *PHYB*) directly interact with *PIFs*, which trigger phytohormone synthesis and signal transduction to regulate plant growth and development. Here our results confirmed that three *PIF* genes (*PIF2, PIF5* and *PIF6*) and several SAS-related auxins (*i.e., IAA, GH3* and *SAUR*) and brassinosteroids (*i.e., BZR1*) genes were significantly up-regulated in the Arabidopsis leaf tissue under the two dim-light conditions (Fig. 5 and Table S8). In contrast, none of the *PIF* genes exhibited significant up-regulation in the two dim-light conditions ginseng leaf tissue (Fig. 5 and Table S6). Our WGCNA results also confirmed that although the co-expressed genes of *PHYB* were functional enriched in similar GO terms between Arabidopsis and ginseng, the PIFs were only identified in Arabidopsis co-expressed network (Fig. S7 and Table S11-S12). In addition, although the blue light receptors *CRY1/2* and hub gene *COP1* showed no significant up-regulation in both the Arabidopsis and ginseng under dim-light conditions. Regulation patterns of these SAS-related genes suggest that the shade avoidance regulatory network was only activated in Arabidopsis leaf tissue under dim-light conditions. It may explain why neither the one-year nor the four-year-ginseng plants did not show any obvious SAS in the phenotypic analysis. It is notable that several genes participating in the biosynthesis and transduction of defense-related jasmonic acid (JA) (*i.e*., *JAZ* and *MYB2*) were significantly up-regulated in both the two species. For example, 11 of the 12 *JAZ* genes are activated in ginseng leaf tissue (Table S6). However, only two of the ten *JAZ* genes exhibited up-regulation in Arabidopsis leaf tissue under the three light conditions (Table S8). These findings together suggest that gene regulation of photomorphogenesis in ginseng differs from Arabidopsis under dim-light conditions (Table S6-9).

**Figure 5.**
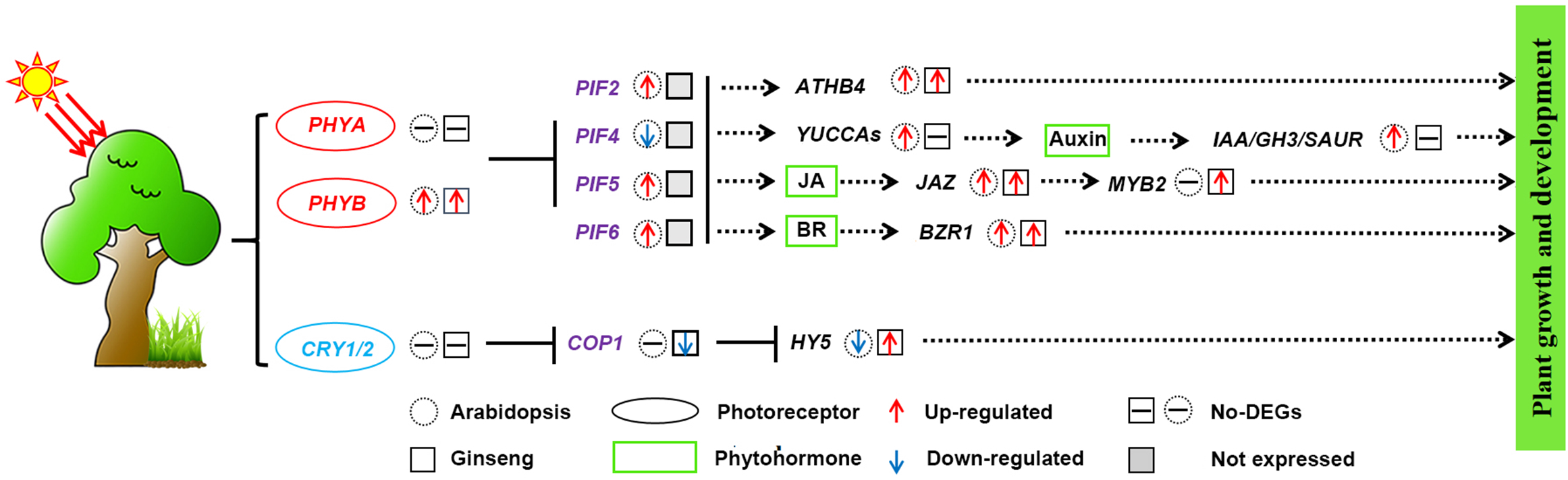
Gene expression patterns of the photomorphogenesis regulatory network in ginseng leaf tissue. From left to right represents the photomorphogenesis regulation process that starts from light receptor (red and blue) to hub genes (purple) and downstream phytochromes-related genes (black). The green rectangles represent three phytohormones involved in shade avoidance and plant defense. The ellipse shape represents red, far-red and blue light receptors. The dashed circle and solid square shapes represent the Arabidopsis and ginseng, respectively. The square shape filled by grey color indicates the deletion of *PIFs* in ginseng genome. Red and blue arrows represent up- and down-regulation of the gene transcription under dim-light compared sunlight condition. Square with black horizontal line represents no differentially expressed gene. T-bar and dashed arrows represent the inhibition and promotion relations between the genes. JA: jasmonic acid; BR: brassinosteroids.

### Genetic bases underlying the shade-tolerance phenotype of ginseng

The above phenotypic comparisons indicate that ginseng plants are more tolerant to shade environment compared to Arabidopsis (see Fig. 1 and 2). We then examined the genetic bases underlying the shade-tolerance phenotype of ginseng. As expected, a total of 28 (43.75%) photosynthesis-related genes possessed more than two-times of copy number in the allotetraploid ginseng genome compared to the diploid Arabidopsis, particularly those of involved in carbon fixation process (*i.e*., *GAPDH*, *PGK* and *RCA*) (Fig. 6A). In contrast, only 15 (23.43%) of these genes exhibited less than two times of copy numbers in ginseng genome relative to Arabidopsis. Multiple gene copies together with high gene transcription level of these photosynthesis-related genes are potentially associated with the high individual performance of ginseng plants under dim-light conditions.

**Figure 6.**
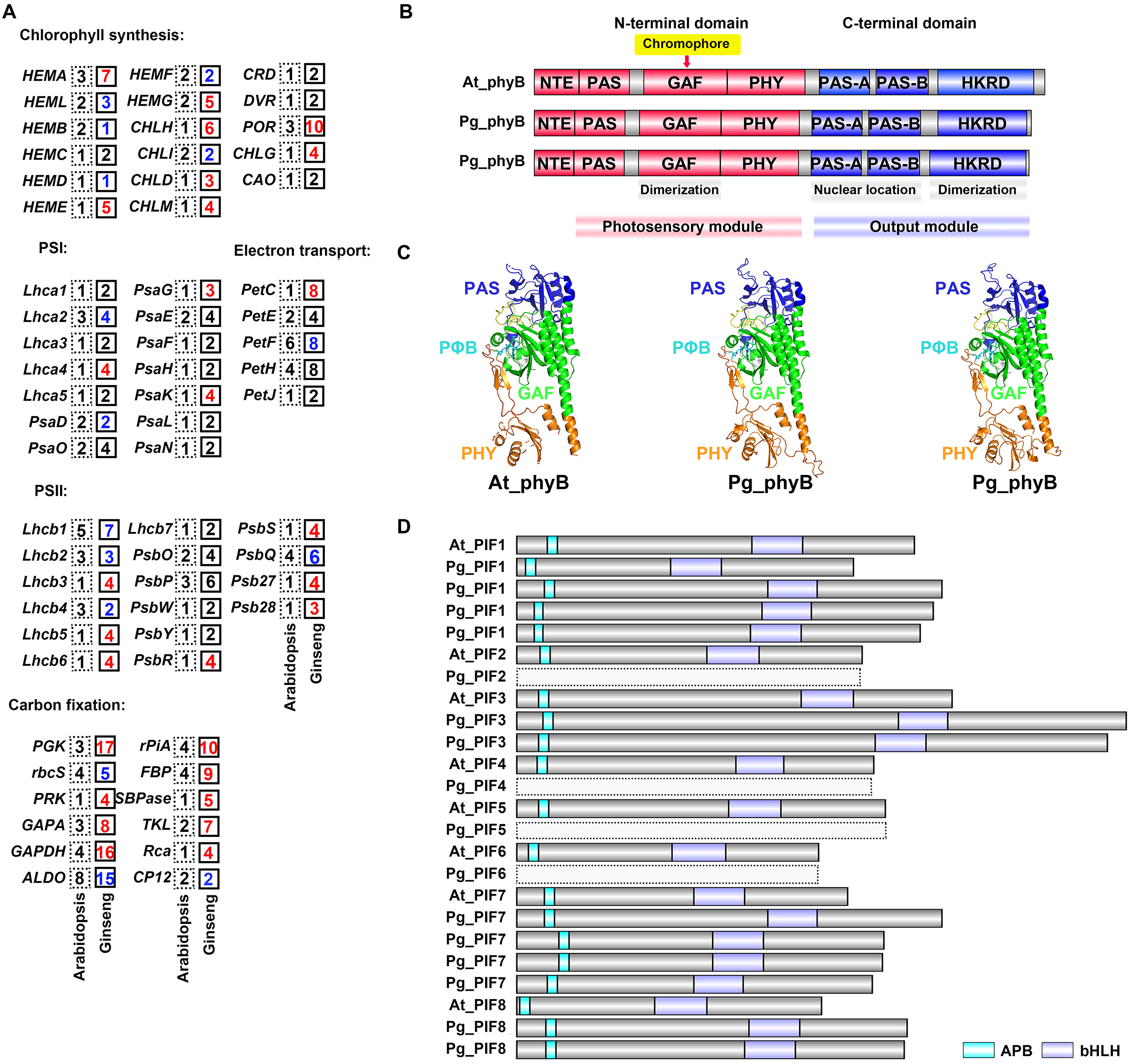
Functional analyses of the photosynthesis- and photomorphogenesis-related genes in ginseng and Arabidopsis. (A) Copy numbers of the photosynthesis-related genes in ginseng and Arabidopsis genomes. Dashed and solid squares indicate the Arabidopsis and ginseng, respectively. Numbers in the square are the copy numbers of each gene. Red, black and blue colors indicate >2-fold higher, equal and <2-fold lower copy numbers in ginseng genome compared to Arabidopsis. (B) Gene structure of *PHYB* in Arabidopsis and ginseng, respectively. NTE, N-terminal extension; PAS, Per (period circadian protein)-Arnt (Ah receptor nuclear translocator protein)-Sim (single-minded protein); GAF, cGMP-stimulated phosphodiesterase, Anabaena adenylate cyclases and *Escherichia coli* FhlA; PHY, phytochrome; PRD, PAS-related domain; HKRD, histidine kinase–related domain. The chromophore is attached to a conserved cysteine residue in the GAF domain. (C) Simulated three-dimension structure of the photosensory module (PSM) of Arabidopsis (At_phyB) and ginseng (Pg_phyB). The PΦB, PAS, GAF, and PHY domains are colored in cyan, blue, green, and orange, respectively. (D) Gene structure of the *PIFs* in Arabidopsis and ginseng. Gene names with the initial “At” and “Pg” represent Arabidopsis and ginseng, respectively. The helix–loop–helix (bHLH) domain is responsible for the dimerization and DNA-binding of PIF protein. Active phytochrome-binding (APB) domain is the binding sites with phyB. The absent of the four *PIF* members (PIF2, PIF4, PIF5 and PIF6) in ginseng was is represented by dashed boxes.

We also performed functional comparisons of *PHYB* and *PIFs*, the key regulatory factors of photomorphogenesis network, between Arabidopsis and ginseng. Our functional analyses of the *PHYB* gene revealed that both the two orthologous copies of ginseng showed similar gene structure with Arabidopsis (Fig. 6B). Further three-dimensional structural simulation of the phyB photosensing module also confirmed normal molecular function in both the Arabidopsis and ginseng (Fig. 6C). In contrast, our functional analyses of the *PIFs* revealed that the four *PIF* members (*PIF2*, *PIF4*, *PIF5*, and *PIF6*) up-regulated in Arabidopsis under dim-light conditions were absent in ginseng genome (Fig. 6D and Fig. S8). The remaining four *PIF* members (*PIF1*, *PIF3*, *PIF7*, and *PIF8*), which showed no differential expression in ginseng, all possessed functional APB motif to interact with *PHYB*. These results suggest that loss-function and non-activation of the PIFs are potentially responsible for the low level of phenotypic plasticity of ginseng plants under different light conditions.

## Discussion

### Low degree of phenotypic plasticity of ginseng plants under distinct light conditions

Plants are autotrophic species that convert solar energy into a chemical form stored in organic compounds (Freudenthal *et al*., 2020). In natural habitats, however, both the intensity and composition of light are highly dynamic on temporal and spatial scales (Hudson *et al*., 2017). Plant species have thus evolved shade-avoidance and shade-tolerance strategies to cope with variable light conditions. Phenotypic responses of heliophytic plants to changing light conditions have been well-documented in model species such as Arabidopsis (Hornitschek *et al*., 2012; Inoue *et al*., 2016). However, it has remained largely unclear of how sciophytic plants (*i.e*., ginseng) respond to distinct light environments. Here we performed phenotypic comparisons between Arabidopsis and ginseng under simulated sunlight and dim-light conditions. The Arabidopsis plants showed highly variable in flowering time, leaf morphologies and plant performance under the different light conditions. However, both the one-year and four-year ginseng plants maintained relatively stable morphological and physiological traits. Under shading stress, the shade-intolerant species prefer to allocate assimilated carbon to the stem and leaf growth (Gommers *et al*., 2017; Pierik and de Wit, 2014). This plant performance confers high phenotypic plasticity on the SAS-related traits (*i.e*., elongation of petioles and stems) to shade-avoidance species to escape from shaded light environments (Valladares and Niinemets, 2008). In contrast, the shade-tolerant species tend to store the fixed carbon energy rather than to maximize plant growth (Gommers *et al*., 2013; Madsen and Owens, 1998). With this reasoning, low phenotypic plasticity of shade-tolerant species may serve as a survival strategy to maintain the balance of survival-growth-reproduction under changing light conditions.

It is notable that ginseng plants have evolved some specific shade-tolerance phenotypes to adapt to shaded habitat. For morphological traits, the ginseng seed is a hypogeal germination type that contains large endosperm to store a considerable amount of carbohydrates. In addition, both the one-year and four-year ginseng plants showed tendency of increased specific leaf area under dim-light conditions. Yet, hypogeal germination is also found in the other shade-intolerant species (Vogel, 1980). Similarly, the increase in specific leaf area is also not a common phenotype in all shade-tolerant plants (Poorter *et al*., 2019). In ginseng, however, the two advantageous phenotypes may not only provide resource of energy during seed gemination but also increase the capacity to capture limited light under shaded habitat. For physiological traits, both the one-year and four-year ginseng plants possessed relatively lower light saturation point (300-400 μmol·m^-2^·s^-1^) compared to Arabidopsis (700-800 μmol·m^-2^·s^-1^) and other heliophytic plant species (600-1000 μmol·m^-2^·s^-1^) (Boonchai *et al*., 2018; Li *et al*., 2018). In particular, we showed that both one-year and four-year ginseng plants exhibited significantly higher Fv/Fm values under dim-light compared to sunlight condition. In contrast, an opposite variation pattern of the Fv/Fm was observed in Arabidopsis. Overall, our phenotypic analyses suggest that high photosynthetic efficiency together with low phenotypic plasticity offer ginseng plants high adaptability to survive in shaded habitat.

### Regulatory network underlying the low phenotypic plasticity in ginseng

Molecular response and regulatory network underlying the SAS have been well-documented in Arabidopsis (Pedmale *et al*., 2016; Sullivan and Deng, 2003; Xie *et al*., 2017). High plasticity of SAS-related traits observed in Arabidopsis can be explained by the rapid responses of red/far-red light and/or blue-light photoreceptors under shaded light condition (Inoue *et al*., 2016; Paik *et al*., 2019). Then, the activated photoreceptor directly interacts with *PIFs* or other hub genes (*i.e*., *COP1*) to promote downstream synthesis of auxin and brassinosteroids, which subsequently regulate plant growth and development to escape the shaded environment (Kozuka *et al*., 2010; Yang *et al*., 2018). Here we aimed to address whether the low phenotypic plasticity of shade-tolerant species was due to the distinct regulatory mechanism of photomorphogenesis network. Compared to Arabidopsis, our comparisons of time-course expression profiling indicate that while ginseng plants also showed rapid molecular responses to distinct light conditions, key regulatory genes involved in photomorphogenesis network did not show dramatic expression-level differences between the sunlight and dim-light conditions. For example, although both the light quality (red:far red ratio) and quantity (red and blue light) varied among the simulated sunlight and dim-light conditions, only the *PHYB* gene showed significant up-regulation under the dim-light condition in ginseng leaf tissue. In Arabidopsis, the activated *PHYB* gene interacts with three *PIFs* (i.e., *PIF4*, *PIF5* and *PIF7*) to promote the SAS under shaded light environments (Hornitschek *et al*., 2012; Yang *et al*., 2018). Our study also confirmed that three *PIF* members (*PIF2*, *PIF5* and *PIF6*) are significantly up-regulated in Arabidopsis under dim-light conditions. However, we did not observe up-regulation of the *PIFs* in ginseng leaf tissue under dim-light conditions. More importantly, four *PIF* members (*PIF2*, *PIF4*, *PIF5* and *PIF6*) are absent in the ginseng genome. Further structural comparisons also confirmed that both the *PHYB* and retained *PIFs* possess normal functions in ginseng. These attributes support the previously proposed PIF-regulation hypothesis that differential control of the *PIFs* can preclude SAS in shade-tolerant species (Gommers *et al*., 2013). In addition, some other antagonist factors (*i.e., PHYA*) have also been proposed to explain the form of shade-tolerance phenotypes (Valladares and Niinemets, 2008). Here we showed that although some candidate genes (*i.e., HY5*) may potentially act as antagonistic factors in the photomorphogenesis regulatory network. However, the downstream target genes involved in phytohormone synthesis are not significantly up-regulated under dim-light conditions in ginseng leaf tissue. These findings together suggest that inactivation of the regulatory photomorphogenesis network under dim-light conditions is a potential mechanism in maintaining low phenotypic plasticity of ginseng in shaded habitat.

### Candidate genes associated with the carbon gain

The relative importance of traits associated with survival and growth under shaded habitat have long been debated (Poorter *et al*., 2019). The carbon gain hypothesis holds that the top priority of a shade-tolerant species is to maximize the photosynthetic carbon gain under low light environment (Givnish, 1988). Here our comparisons of the leaf transcriptome expression profiling between Arabidopsis and ginseng revealed distinct transcriptional regulations at these genes involved in pigment synthesis, light capture and electron transport. For example, phenotypic analysis revealed higher chlorophyll a/b content in both the two species under dim-light compared to sunlight condition. Consistent with this observation, genes involved in the chlorophyll a/b synthesis (*i.e., HEMA* and *CAO*) were relatively up-regulated under dim-light conditions in both the two species. It is notable that several genes related to light capture (*i.e., LHC-I* and *LHC-II*) and electron transport (*i.e., PetE* and *PetF*) are specifically up-regulated under dim-light conditions in ginseng leaf tissue. This may explain why the ginseng plants exhibited higher photosynthetic efficiency (*i.e*., Fv/Fm value) under dim-light conditions. In Arabidopsis, while these plants grown under sunlight exhibited higher Fv/Fm value, majority of these genes are either down-regulation under sunlight condition (*i.e., Lhca1* and *Lhcb1*) or no differential expressed among the three light conditions (*i.e., PsbI* and *PsbK*). Likewise, core-genes of both the PS-I (*i.e., PsaA* and *PsaC*) and PS-II (*i.e., PsbA* and *PsbC*) were specifically up-regulated under sunlight condition in ginseng leaf tissue, possibly due to the repair of PS-I and PS-II complex under sunlight condition. In contrast, no similar regulation pattern was observed at these PSI and PSII genes in Arabidopsis leaf tissue. It is possible that phenotyping of the two species were estimated using these plants grown under three light conditions for three weeks. However, plants for transcriptomic analyses were grown under their simulated natural environments and then transferred to the three distinct light conditions. Differences in transcriptomic and phenotypic responses suggest higher adaptability of ginseng plants to changing light conditions compared to the Arabidopsis.

Our phenotypic comparisons also revealed relatively smaller plant size of Arabidopsis under dim-light compared to sunlight condition. However, the one-year and four-year ginseng plants showed very similar plant size among the three light conditions. This observation together with leaf morphologies and anatomical structure suggested that ginseng plants may possess high carbon fixation efficiency under dim-light conditions. Consistent with this assumption, we observed relatively higher expression level of the carbon fixation genes (*i.e., rbcL* and *GAPA*) in ginseng leaf tissue under dim-light compared to the sunlight condition. In addition, three genes (*RCA* and *CP12*) involved in the activation of carbon fixation genes were also up-regulated in dim-light conditions, especially for the deep-shade light condition. In contrast, most of these carbon fixation genes did not show dim-light specific up-regulation in Arabidopsis leaf tissue. More importantly, majority of these genes have more copy numbers in ginseng genome than the Arabidopsis. These attributes indicate that these candidate genes may increase the acquisition of carbon in the low light condition of shaded habitat.

### Plant performance under shading stress

Our phenotypic comparisons between the two species support the carbon gain hypothesis that ginseng has developed specialized morphological and physiological traits to improve light capture and use efficiency under dim-light conditions. Nevertheless, compared to the one-year seedling, the four-year ginseng plant showed different variation patterns in leaf morphologies and chlorophyll content under deep-shade condition. It may support the stress tolerance hypothesis that shade-tolerant species prefer to allocate energy and resource to survival-related traits (*i.e*., herbivores and pathogens resistance) under shade stress rather than the maximization of carbon gain (Kitajima, 1994; Poorter *et al*., 2019). In Arabidopsis, it has been shown that lower stress tolerance was observed under simulated shade relative to the sunlight condition (Liu *et al*., 2019). Indeed, genes involved in JA synthesis tend to show down-regulated under shaded light conditions (Fernandez-Milmanda *et al*., 2020). Here, our transcriptomic analyses also confirmed that none of the *JAZ* genes are specifically up-regulated under dim-light condition. In contrast, we identified more up-regulation JAZ genes in ginseng leaf tissue under dim-light conditions. Previous studies have clearly documented that the JA-mediated signaling plays an important role in improving the plant’s defense capabilities (Zander *et al*., 2020). Under this hypothesis, the activation of defense-related phytohormone JA, together with distinct plant performance, under deep-shade condition suggest that the ginseng plants may employ an alternative strategy to balance the survival-growth trade-off under deep-shade light condition.

## Abbreviations

SAS: shade-avoidance syndrome
*PIFs*: phytochrome-interacting factors

## Data availability

All data generated in this study were submitted to GenBank under the Bioproject numbers PRJNA742834 and PRJNA775987.

## Acknowledgments

We thank Prof. Kenneth M. Olsen at Washington University in St. Louis for his constructive comments and edits on previous version of this manuscript. This work was financially supported by National Natural Science Foundation of China (31970235), China Postdoctoral Science Foundation Grant (2018M630400), Shanghai Pujiang Program (19PJ1401500) and Start-up funding at Fudan University (JIH1322105).

## Author contributions

L.F.L. conceived this project; Y.X.Z., Z.H.W. and L.F.L. designed the experiment; Y.X.Z., Y.Q.N., X.F.W., Z.H.W., M.L.W., and Z.H.W. carried out experiments and analyzed the data; Y. X. Z., Y. Q. N., X. F. W., Z. H. W., M. L. W., J. Y., Z. P. S., Y. G. W., W. J. Z and L. F. L interpreted the data and participated in discussion; Y.X.Z., J.Y., Z.P.S., Y.G.W., W.J.Z and L.F.L. wrote the manuscript. All authors contributed to this work gave final approval for publication.

## Supporting Information

**Table S1.** The exact light intensity in natural habitats.

**Table S2.** Descriptive statistics of phenotypic and physiological data. All the data were shown in Figure 2.

**Table S3.** Maximum quantum yield of PS II (Fv/Fm) of Arabidopsis and ginseng under sunlight and dim-light conditions. All the data were visualized in Figure 1D.

**Table S4.** Statistical data for measuring light response curves of Arabidopsis and ginseng. All the data were visualized in Figure 2A.

**Table S5.** Pathways with significant enrichment of DEGs under sunlight and dim-light conditions. All the data were visualized in Figure 3D.

**Table S6.** GO enrichment analysis of DEGs under sunlight and dim-light conditions in Arabidopsis and ginseng.

**Table S7.** Fold changes of DEGs between the treatment (0.25-12 h) and control (0 h) time points under the same conditions in ginseng.

**Table S8.** Fold changes of DEGs at the same treatment time point between the dim-light and sunlight conditions in ginseng.

**Table S9**. Fold changes of DEGs between the treatment (1-6 h) and control (0 h) time points under the same conditions in Arabidopsis.

**Table S10**. Fold changes of DEGs at the same treatment time point between the dim-light and sunlight conditions in Arabidopsis.

**Table S11**. Candidate genes that are co-expressed with *PHYB* in Arabidopsis and Ginseng, respectively.

**Table S12**. Functionally enriched GO terms of the candidate genes that showed co-expressed with *PHYB* in Arabidopsis and Ginseng, respectively.

**Figure S1.** Paraffin sections of Arabidopsis and ginseng under sunlight and dim-light conditions. Bar represents 50 μm.

**Figure S2.** Visualizing transcriptome data based on expression diversity (*Hj*) and specificity (*δj*) at the subgenome level. Each symbol represents a sampling time point.

**Figure S3.** Clustering heatmap of the total gene of ginseng (A) and Arabidopsis (B) under the three light conditions. Different colors represented the expression-level difference of the samples. Numbers on the right of the bar indicate the level of expression-level difference. Colors from light to dark represent the similarity of gene expression pattern.

**Figure S4.** Clustering heatmap of the DEGs under sunlight and dim-light conditions in ginseng (A) and Arabidopsis (B). Color scheme indicates the up (red) and down (blue) of the genes between the treatment (0.25-12 h) and control (0 h) time point.

**Figure S5.** Venn plot of the number of DEGs at the subgenomic level. (A) Subgenome A (B) Subgenome B. Numbers within each circle were obtained from the comparisons between the control and six treatment time points at the subgenomic level.

**Figure S6**. Expression pattern of the photosynthesis-related genes in Arabidopsis under sunlight and dim-light conditions. (A) Differentially expressed genes (DEGs) involved in chlorophyll synthesis pathway. (B) DEGs involved in PSI, PSII and electron transport chain. (C) DEGs involved in carbon fixation. Gene names marked in black and green color represent nuclear and chloroplast genes, respectively. Color scheme indicates the up (red) and down (blue) of the genes between the treatment (1, 3, and 6 h) and control (0 h) time point. PC: Plastocyanin; PQ: Plastoquinone; Fd: Ferredoxin; FNR: Ferredoxin-NADP+-oxidoreductase; Cyt b6/f: Cytochrome b6/f complex; PSI: Photosystem I; PSII: Photosystem II.

**Figure S7**. Identification of the *PHYB* co-expression genes and its molecular functions in ginseng and Arabidopsis. Genes that showed co-expression with *PHYB* in Arabidopsis (A) and ginseng (B) under dim-light conditions. (C) functionally enriched GO terms of the genes co-expressed genes in Arabidopsis and Ginseng.

**Figure S8**. Protein sequence alignment of the APB and bHLH motifs in PIFs. The helixloop–helix (bHLH) domain is responsible for dimerization and DNA-binding of the PIF protein. active phytochrome-binding (APB) domain is the binding sites with phyB.

