## Supplementary material for "Phenotypic and transcriptomic responses of the sun- and shade-loving plants to sunlight and dim-light conditions": Fig. S

### Slide 1
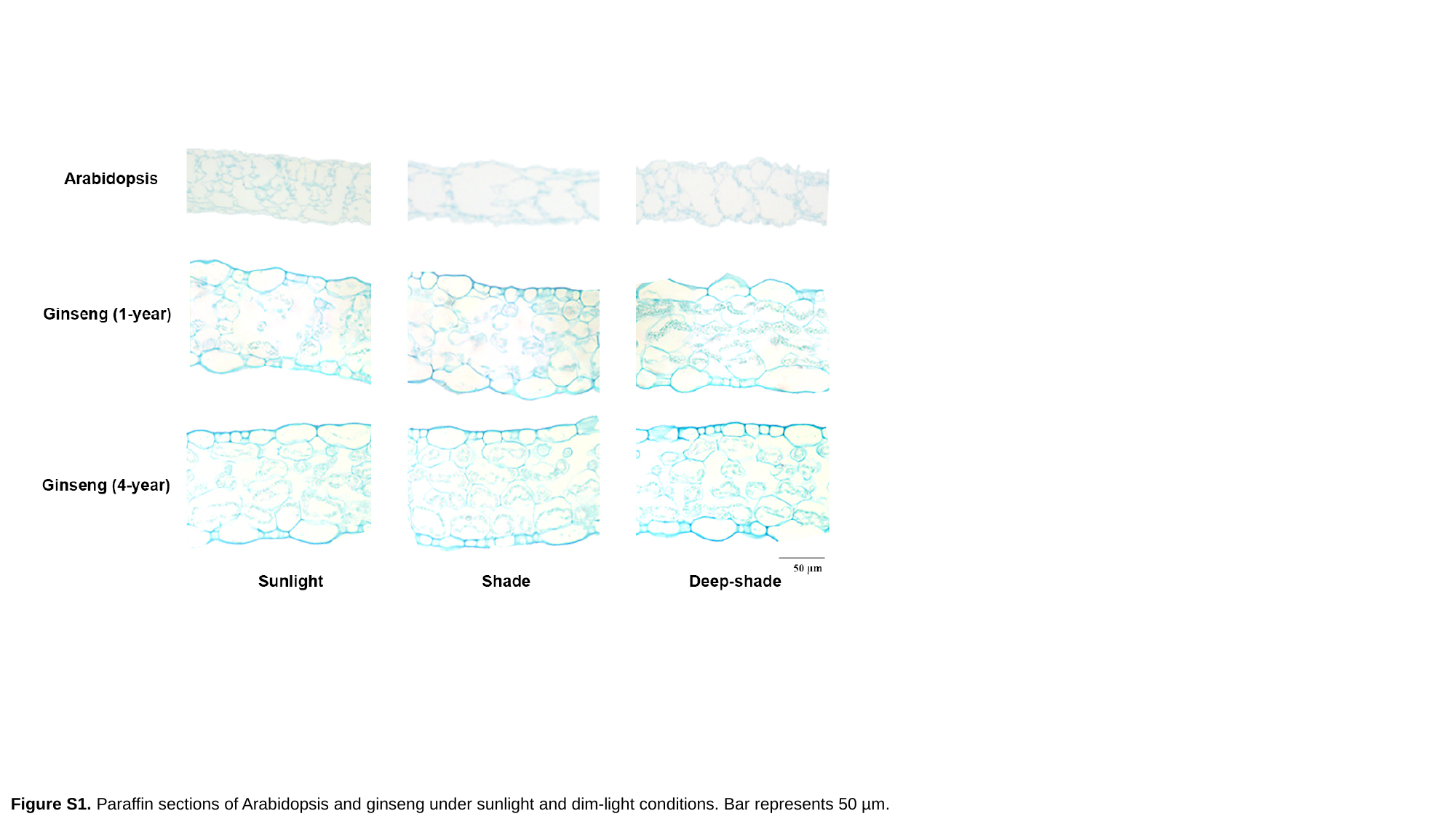

Figure S1. Paraffin sections of Arabidopsis and ginseng under sunlight and dim-light conditions. Bar represents 50 µm.

### Slide 2
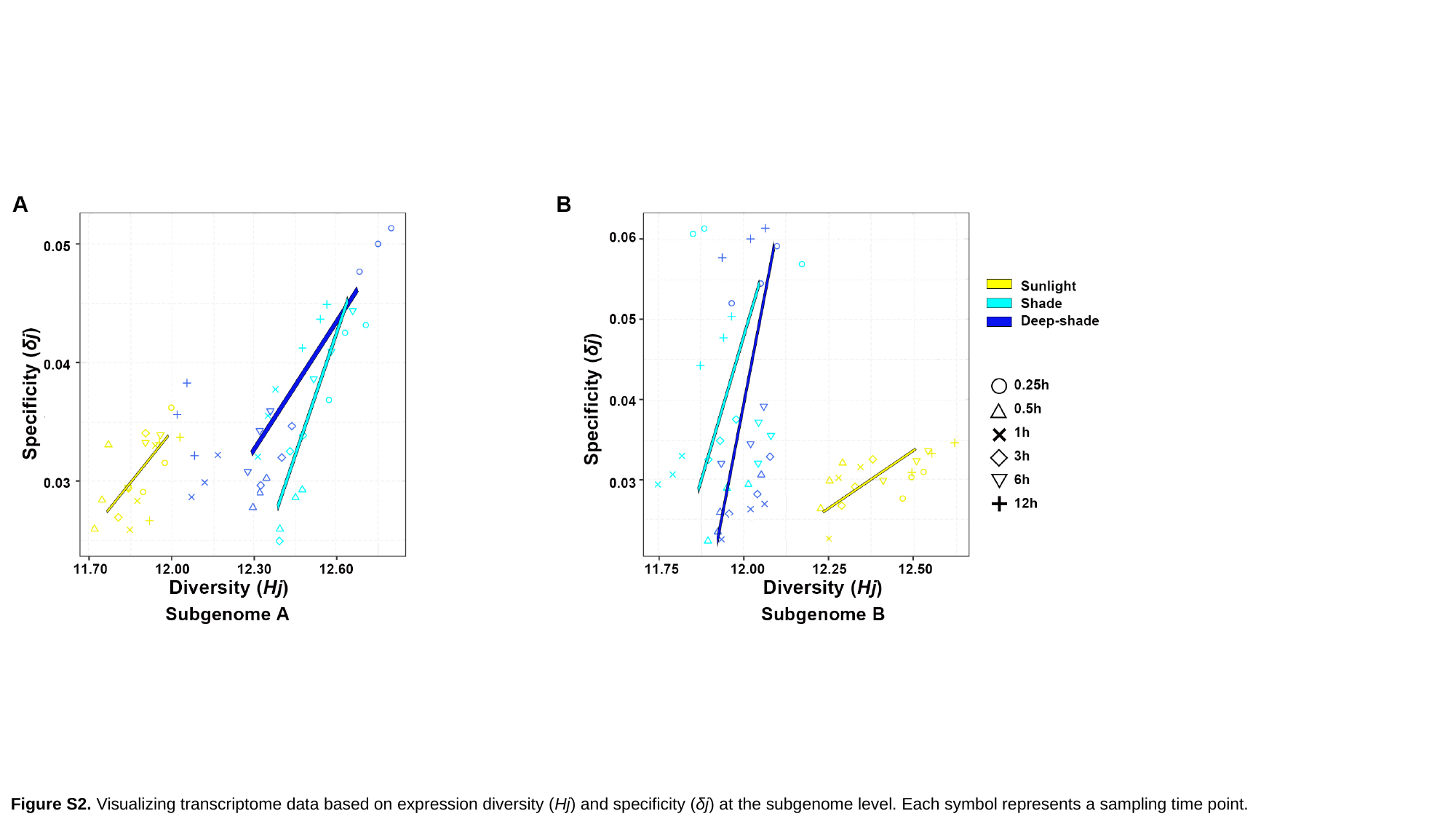

Figure S2. Visualizing transcriptome data based on expression diversity (Hj) and specificity (δj) at the subgenome level. Each symbol represents a sampling time point.

### Slide 3
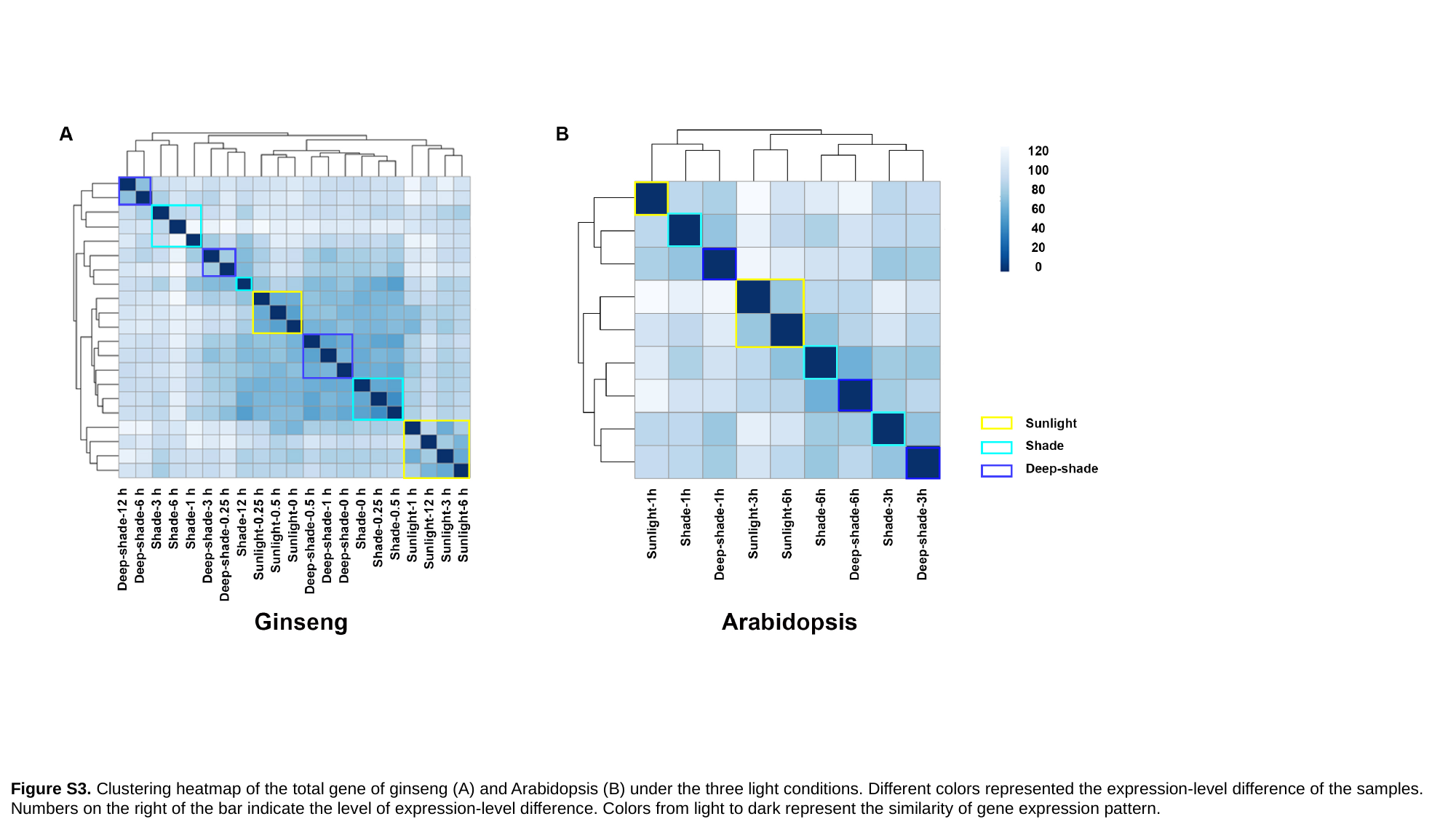

Figure S3. Clustering heatmap of the total gene of ginseng (A) and Arabidopsis (B) under the three light conditions. Different colors represented the expression-level difference of the samples. Numbers on the right of the bar indicate the level of expression-level difference. Colors from light to dark represent the similarity of gene expression pattern.

### Slide 4
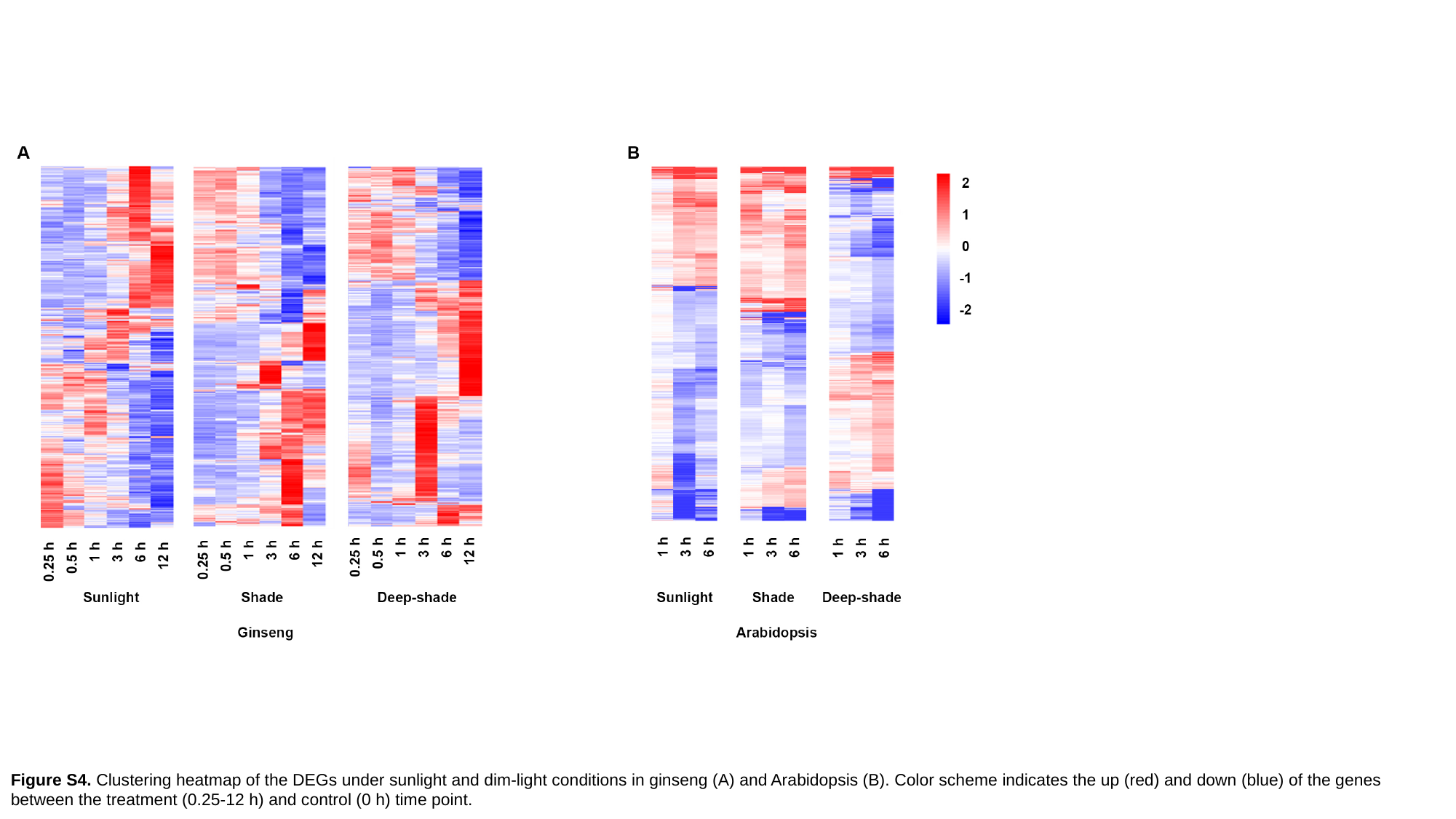

Figure S4. Clustering heatmap of the DEGs under sunlight and dim-light conditions in ginseng (A) and Arabidopsis (B). Color scheme indicates the up (red) and down (blue) of the genes between the treatment (0.25-12 h) and control (0 h) time point.

### Slide 5
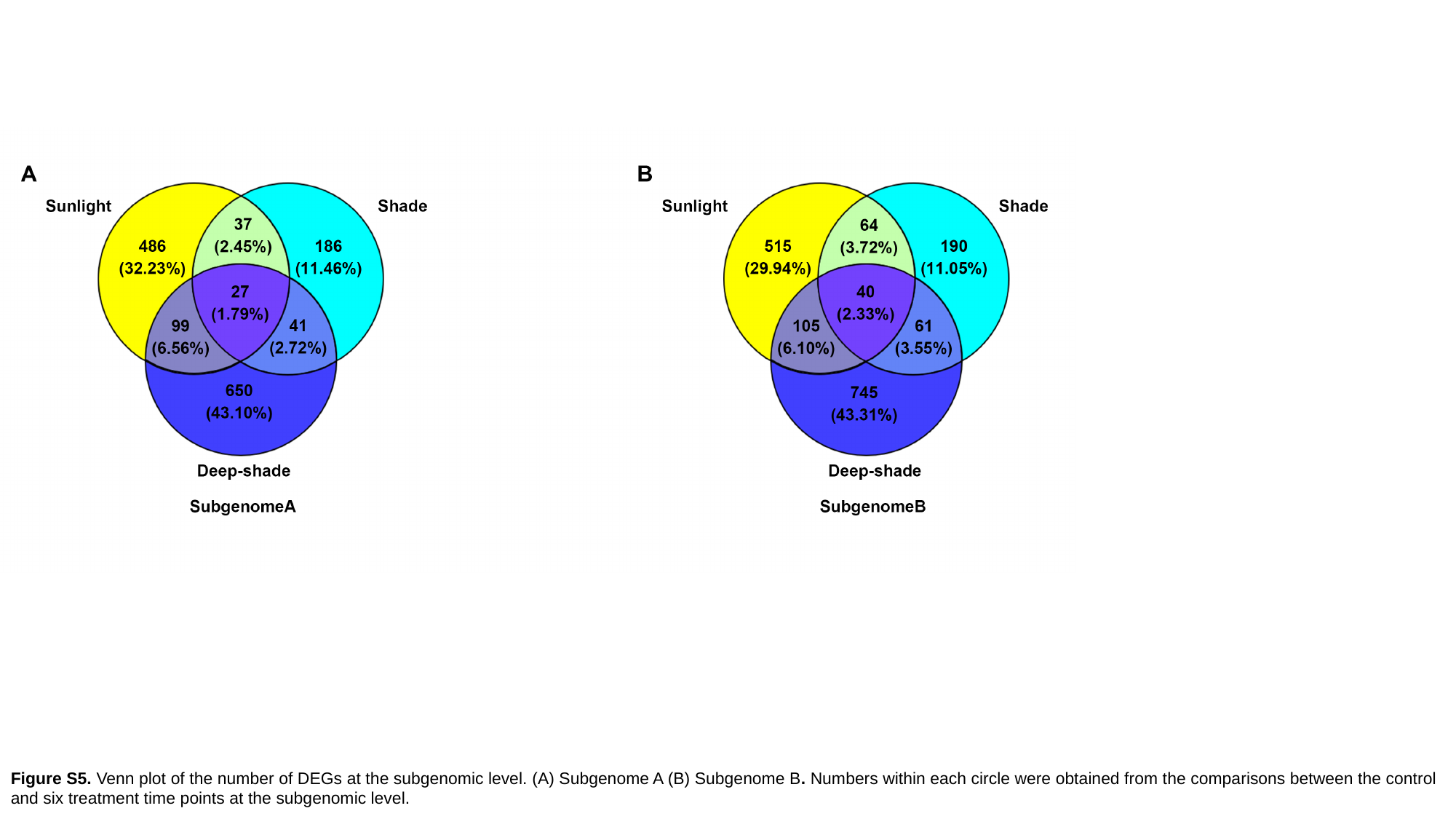

Figure S5. Venn plot of the number of DEGs at the subgenomic level. (A) Subgenome A (B) Subgenome B. Numbers within each circle were obtained from the comparisons between the control and six treatment time points at the subgenomic level.

### Slide 6
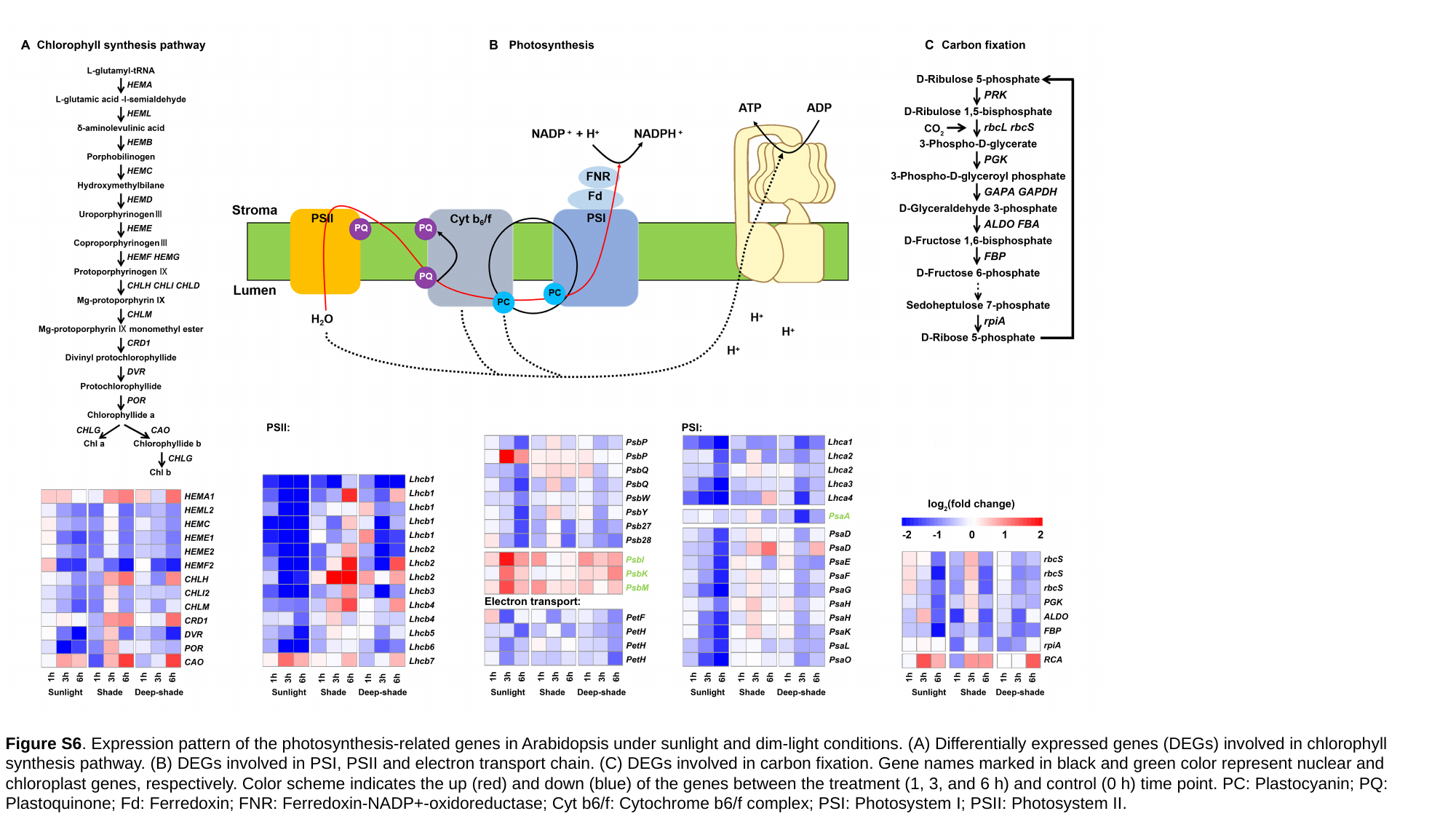

Figure S6. Expression pattern of the photosynthesis-related genes in Arabidopsis under sunlight and dim-light conditions. (A) Differentially expressed genes (DEGs) involved in chlorophyll synthesis pathway. (B) DEGs involved in PSI, PSII and electron transport chain. (C) DEGs involved in carbon fixation. Gene names marked in black and green color represent nuclear and chloroplast genes, respectively. Color scheme indicates the up (red) and down (blue) of the genes between the treatment (1, 3, and 6 h) and control (0 h) time point. PC: Plastocyanin; PQ: Plastoquinone; Fd: Ferredoxin; FNR: Ferredoxin-NADP+-oxidoreductase; Cyt b6/f: Cytochrome b6/f complex; PSI: Photosystem I; PSII: Photosystem II.

### Slide 7
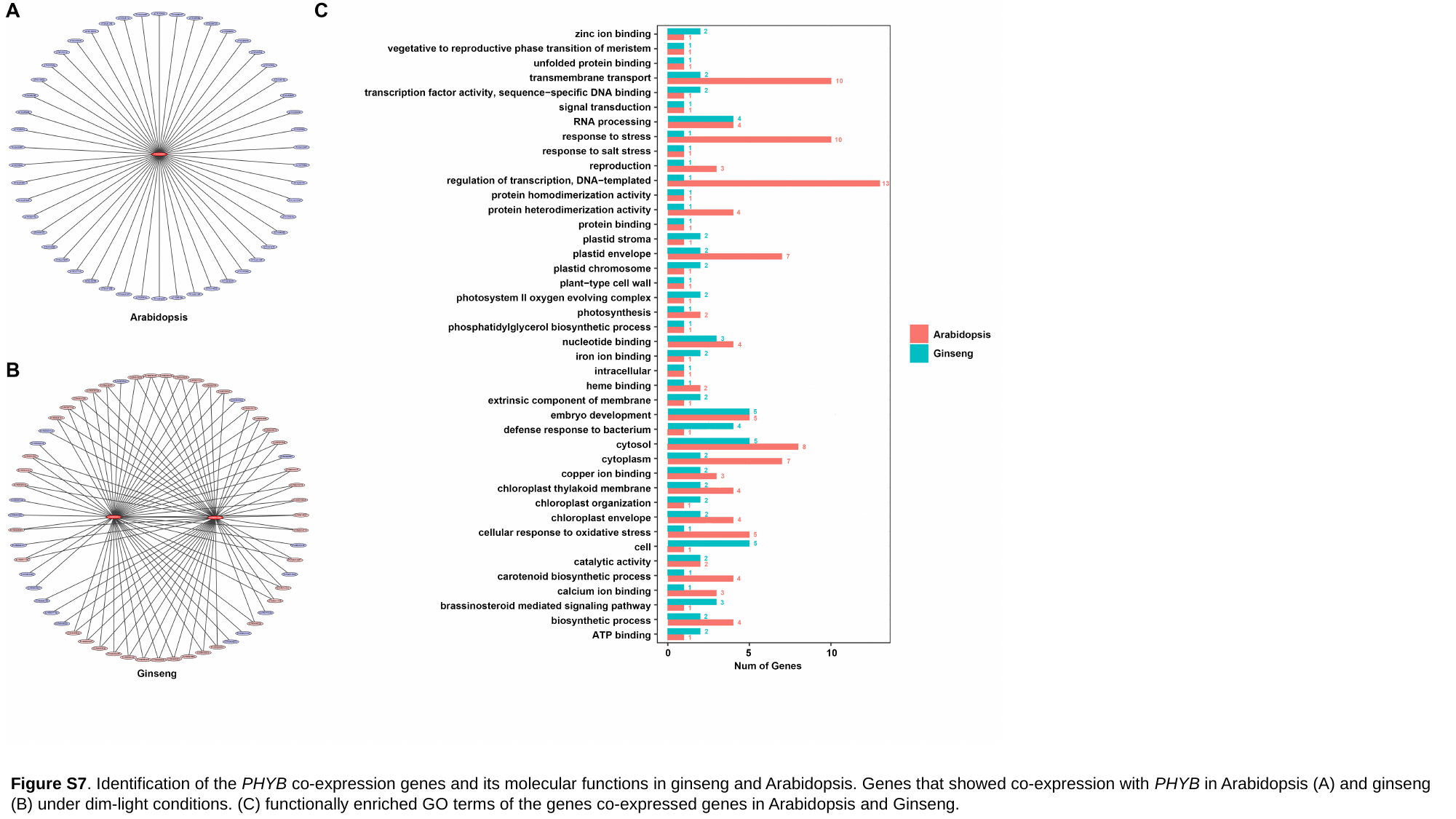

Figure S7. Identification of the PHYB co-expression genes and its molecular functions in ginseng and Arabidopsis. Genes that showed co-expression with PHYB in Arabidopsis (A) and ginseng (B) under dim-light conditions. (C) functionally enriched GO terms of the genes co-expressed genes in Arabidopsis and Ginseng.

### Slide 8
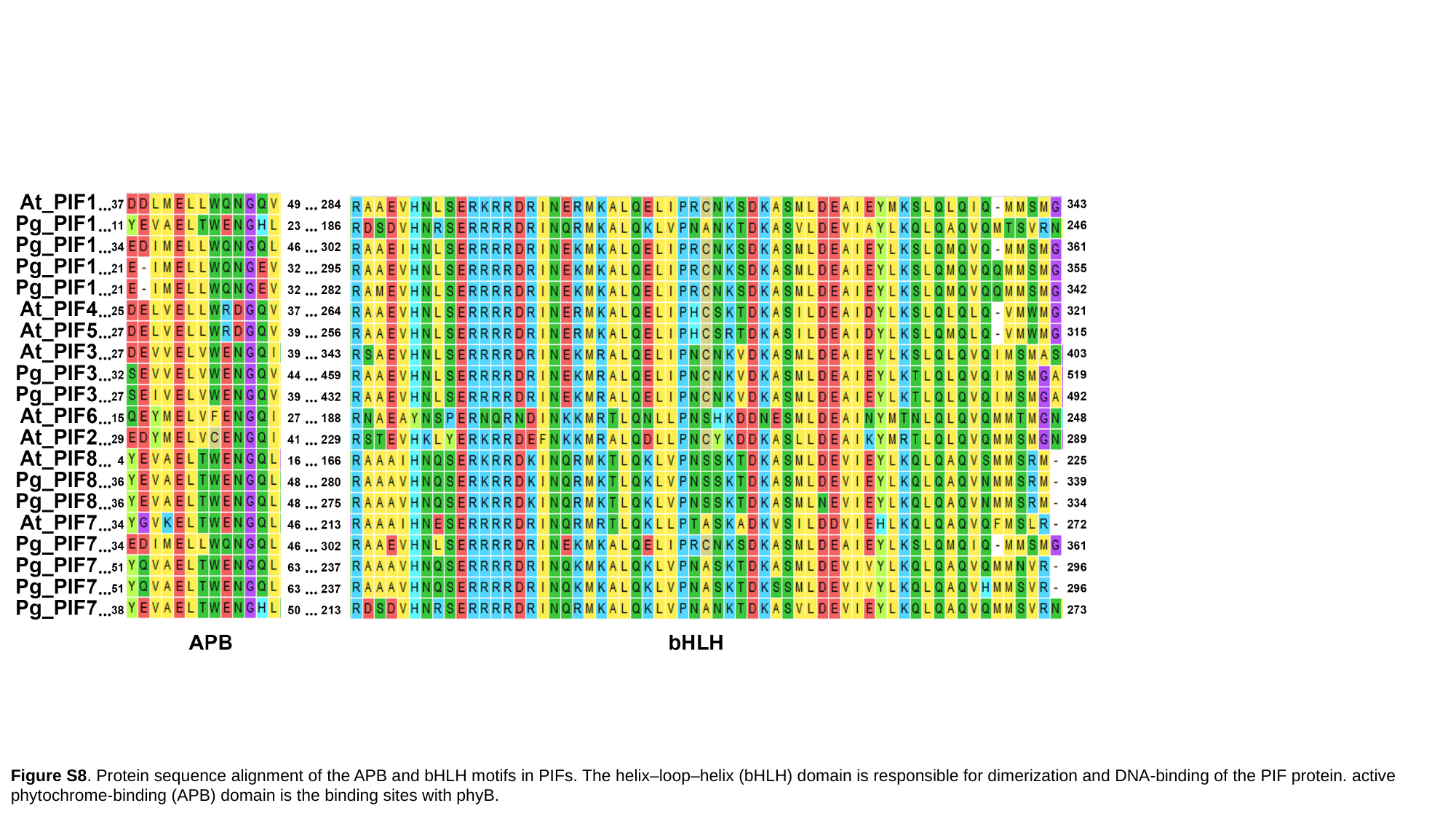

Figure S8. Protein sequence alignment of the APB and bHLH motifs in PIFs. The helix–loop–helix (bHLH) domain is responsible for dimerization and DNA-binding of the PIF protein. active phytochrome-binding (APB) domain is the binding sites with phyB.
